# Scarcity of flowering plants on *Sedum* roofs limits pollinator diversity

**DOI:** 10.1101/2025.09.14.676189

**Authors:** Eva Drukker, Marco Tanis, Leon Marshall, Casper W. Quist, M. Eric Schranz, Nina E. Fatouros

## Abstract

Green roofs can provide suitable habitats for pollinating insects in urban areas. Pollinators are a large and diverse group with high economic and ecological relevance to society. Yet their populations are in decline, highlighting the need for new habitats such as green roofs. An understanding of how local and landscape factors of green roofs shape pollinator communities is crucial to optimize the design and development of green roofs.

This study aimed to identify how the surrounding landscape and green roof characteristics influence pollinator diversity and abundance. Pollinators – such as bees, wasps, hoverflies and butterflies - were sampled using pan traps and hand netting on 25 green roofs that were categorized into three types based on functionality and vegetation structure: sedum roofs, nature roofs, and roof gardens. To assess pollinator communities, we examined species richness, abundance, and functional diversity—defined here as the range of ecological roles or traits. Roof characteristics (e.g., flower diversity, microrelief, vegetation cover, roof height, size, and age) and landscape factors (e.g. percentage of surrounding green, distance to surrounding green) were analysed in relation to these diversity metrics.

We show that a continuous supply of flowers throughout the year, flower abundance and the type of green roof significantly influence pollinator diversity and abundance. Sedum roofs supported lower pollinator diversity compared to the other roof types. Furthermore, our models indicated a decrease in pollinator diversity on higher green roofs, and when honeybee hives are present, while the presence of bee hotels increased pollinator abundance.

The study highlights the value of nature roofs and roof gardens, compared to sedum roofs for supporting pollinator diversity. This is linked to the importance of constant forage availability, especially in early spring, which is absent on most sedum roofs. The results from this study offer guidelines to green roof designers on how to design pollinator friendly roofs in urban areas.

**Graphical Abstract:** 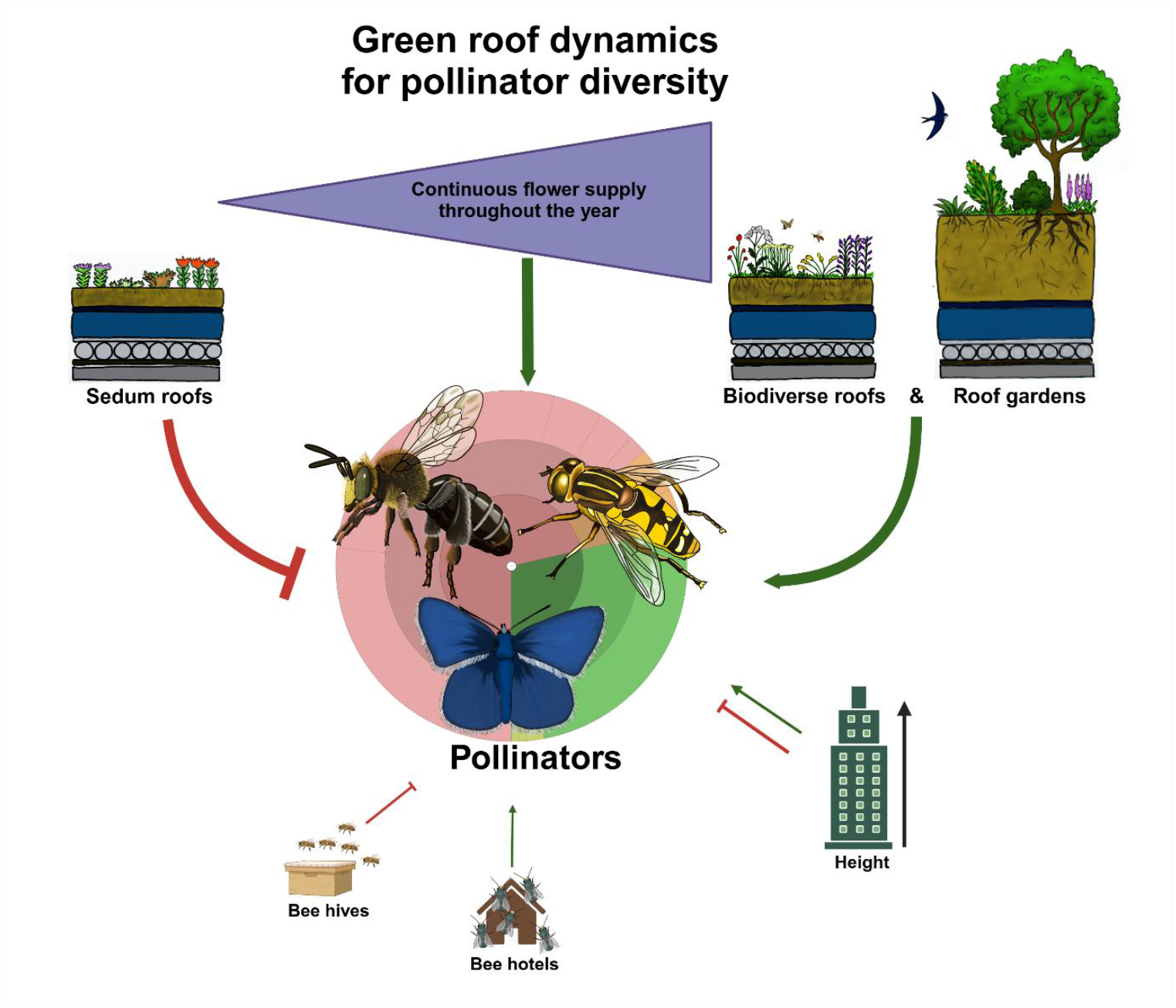

Pollinator species: *Macropis europea* (Hymenoptera: Melittidae), *Helophilus pendulus* (Diptera: Syrhidae) and *Polyommatus icarus* (Lepidoptera: Lycaenidae). All three species were observed on green roofs during the authors’ fieldwork. Image created using biorender.com and personal drawings.

**Highlights:**

- Biodiverse roofs and roof gardens support higher pollinator diversity compared to sedum roofs.
- A continuous supply of flowers throughout the year, along with a higher abundance of flowers, are key factors influencing pollinator diversity and abundance on green roofs.
- Roof height affects the diversity and abundance of different pollinator groups in distinct ways, a novel finding that needs further investigation.
- We provide guidelines for green roof designers, emphasizing the importance of selecting roof types and designs that ensure constant flower availability.

## Introduction

Pollinating insects like wild, native bees, butterflies and hoverflies are an important group to study ecosystem services and the impact of environmental changes on biodiversity (Naeem et al., 2020; Rader et al., 2016; Ricou et al., 2014). Pollinators provide as a source of food for higher trophic levels (e.g. birds and bats) and contribute to plant pollination (Hallmann et al., 2017; Rader et al., 2016). At least 75 percent of food crop species are to some degree dependent on animal pollination, as well as 88 percent of all flowering plants (Ollerton et al., 2011). Their populations are in decline due to agricultural intensification, pesticide use, habitat loss, and climate change (Biesmeijer et al., 2006; Dicks et al., 2021; Hallmann et al., 2017; Sánchez-Bayo & Wyckhuys, 2019). Besides their contribution to ecosystem services, pollinator conservation is important to protect species diversity to ensure long-term ecosystem resilience (Senapathi et al., 2015). Conserving pollinators is thus crucial. Urban green spaces can help mitigate these declines by offering habitat for native and colonizing species (Banaszak-Cibicka et al., 2018; Hall et al., 2017; Wenzel et al., 2020). While urbanization typically reduces pollinator presence (Bates et al., 2011; Clark et al., 2007), some groups like cavity-nesting bees thrive due to increased resources (Cane et al., 2006; Matteson et al., 2008), due to increases in food, nesting and habitat resources (Sexton et al., 2021).

Green roofs represent one potential urban habitat, supporting diverse insect communities (Kadas et al., 2006; Knapp et al., 2019; MacIvor, 2016; Madre et al., 2013; Tonietto et al., 2011). However, conventional sedum roofs offer limited biodiversity due to shallow substrates and uniform vegetation with limited flower and pollen availability (Madre et al., 2013; Williams et al., 2014). In contrast, other roof types, like nature roofs and roof gardens with more diverse vegetation and structure show greater potential for supporting pollinators (Sutton et al., 2015; Williams et al., 2014). These roofs have been claimed to be beneficial for biodiversity compared to sedum roofs, especially for pollinating insects like bees (MacIvor, 2016), though research on their effectiveness remains limited (Coulibaly et al., 2023).

Functional traits offer insight into how pollinators use green roofs for feeding, nesting, and reproduction, and can clarify species’ ecological roles and adaptability (Tilman, 2001). In the context of green roofs, understanding the functional traits of pollinators helps evaluate their ability to use these habitats for key life cycle stages such as feeding, nesting, and reproduction, which is often lacking in green roof research (Coulibaly et al., 2023; Tiago et al., 2024). With these traits, functional richness can be calculated which is a biodiversity index reflecting much of the possible ecological roles that are filled by the species present in the study system (Legras & Loiseau, 2018).

Lastly, the influence of roof design, height, and surrounding landscape on pollinators is not fully understood, especially for butterflies and hoverflies. Bee diversity may decline with roof height (MacIvor, 2016; Tonietto et al., 2011), likely due to their limited mobility (Zurbuchen et al., 2010). However, the effect of height and distance to the surrounding green has not always been found in other pollinator research (Jacobs et al., 2023; Kratschmer et al., 2018).

This study examines pollinator diversity on green roofs—focusing on wild bees, aculeate wasps, hoverflies, butterflies, and others (see Appendix Table A.1) at the species level. Three indices were measured, —namely species richness, abundance, and functional richness. More specifically, we tested (1) how pollinator species richness, functional richness, species composition and abundance on sedum roofs compares to nature roofs and roof gardens, (2) which roof factors increase pollinator diversity and (3) which pollinator traits are typically found on green roofs.

We hypothesize that nature roofs and roof gardens support higher pollinator abundance and diversity than sedum roofs. Because nature roofs and roof gardens tend to support a higher abundance and diversity of flowering plants – and therefore higher pollen availability – compared to sedum roofs, the observed differences in pollinator richness between roof types is likely mediated by floral resources, therefore we expect that flower diversity and abundance will be strong predictors of pollinator community across roof types. We also expect that less mobile, nesting pollinators like solitary bees are more sensitive to roof height and surrounding habitat availability than more mobile, non-nesting pollinators like butterflies and hoverflies. The latter are therefore further grouped and analysed as nesting and non-nesting pollinators. Addressing these questions will enhance our understanding of the relationship between green roof characteristics and pollinator diversity, providing important insights for the design of more pollinator-friendly green roofs.

## Materials and methods

### Study sites

We sampled 25 green roofs spread across multiple urban areas in the Netherlands (Table A.2 for details and appendix Fig. B.1 for map) classifying them into sedum roofs, nature roofs, and roof gardens based on substrate depth, plant composition, and human influence (Bianchini & Hewage, 2012; Madre et al., 2013; Sutton et al., 2015). The nine *sedum* roofs (a.k.a. extensive green roofs) in this study had a substrate layer between 5 to 7 cm. They were primarily composed of the succulent, drought-resistant *Sedum* plants and bryophytes (mosses). The six nature roofs (sometimes referred to as semi-intensive or extensive (Kotze et al., 2020)) had a substrate layer between 6 to 15 cm. They were composed of a combination of *Sedum* plants and herbaceous plants, with herbaceous plants being dominant on more than 20% of its surface and an average of around 35 species per roof. The ten roof gardens had a substrate layer between 15 to 200 cm or more. They were composed of the previous strata as well as shrubs and trees. They require the same (high) maintenance as a normal garden, hence the name (Sutton et al., 2015). Unlike the other roofs, roof gardens were usually accessible to the general public, or residents of the building.

### Flower availability

Four variables of flower availability were recorded: abundance, diversity, species richness and seasonality using sixteen 1 m^2^ plots per roof. Inflorescences were counted per species (see detailed methods in table A.3 (Van Swaay et al., 2011)). Abundance was the total flower units across plots; diversity was calculated using Shannon’s index. Species richness was supplemented with a 10-minute inventory outside plots. Seasonality was measured by applying Shannon’s index to flower abundance across three seasonal surveys (Help et al., 1998), giving a measure of how even the flower supply was throughout the year.

### Other green roof factors

Other factors that could influence pollinator diversity were recorded per roof. Table 1 gives an overview of these factors, the units and methods on how they were recorded.

**Table 1:** Green roof factors recorded per roof, included in study.

| Green roof factors | Unit/categories | Method |
| --- | --- | --- |
| Type of green roof | Sedum – nature – roof garden | Classify in field |
| Roof age | Years |  |
| Roof area (green only) | m<sup>2</sup> | GIS |
| Roof height | m | From Actueel Hoogtebestand Nederland (AHN 2019) |
| Bee hotel | Yes – No | Classify in field |
| Honeybee hives | Number of honeybee hives on roof | Count |
| Flower abundance | Flowering units | Quadrat plots |
| Flower species richness | No. of species | Quadrat plots & 10 min. inventory |
| Flower diversity | Number | Quadrat plots |
| Flower seasonality | Number | Quadrat plots |
| Total of vertical structures | m<sup>2</sup> | Tape measure & satellite imagery |
| Difference in vegetation height | cm | Tape measure |
| Microrelief | None –small amount – large amount | Classify in field |
| Vegetation cover | Percent (See App. C) | Estimate |
| Habitat in surrounding area | m<sup>2</sup>. (See App. F) | GIS |
| Distance to suitable habitat | m (See App. F) | GIS |

### Insect sampling

Pollinators were sampled during three periods in 2019: early spring (8 Apr–15 May), late spring (22 May–4 Jun), and summer (1 Jul–1 Aug). Two methods were used: pan trapping and a timed search with hand nets. On each roof three white, three yellow and two blue pan traps were placed for 24 hours filled with water and a few drops of detergent. Pan traps were made using acrylic-based spray paint on plastic bowls. These colours were chosen because white and yellow pan traps are the most attractive for pollinators, while blue attracts several species not caught by yellow and white (Vrdoljak & Samways, 2012). Additionally, a 20 minute timed search was performed per roof with a butterfly catching net, between 10 a.m. and 5 p.m. Sampling was done under similar weather conditions meaning: no rain, wind speed below 6 Bft, and either a temperature between 12 and 15 degrees Celsius if there was less than 50 percent cloud cover, or a temperature above 15 degrees (adapted from Van Swaay, et al., 2011). Trap duration and weather data were recorded (see table A.4).

### Pollinator identification

Pollinators were identified in the field or preserved (pinned or in alcohol) for later ID. Target groups included wild species (so excluding honeybees) from: Aculeata (bees and wasps, excluding ants), Rhopalocera, Zygaenidae and Macroglossina (Lepidoptera), and several Diptera families (Syrphidae, Stratiomyidae, Conopidae, Bombyliidae). These taxa were chosen for their well-known ecology and pollination relevance (Klecka et al., 2018; Matteson et al., 2008). Pollinators were identified to species level by the authors with the use of identification guides (Bot, S., & Van de Meutter, 2019; Falk, 2017; Paukkunen et al., 2015; Schulten, 2018; Smit, 2013). Some difficult species were identified or checked by experts.

### Pollinator diversity indices

Three metrics were used:

1. Species richness – total number of pollinator species per roof across all sampling rounds.
2. Abundance – total individuals per roof, per sampling round (analysed separately to account for weather).
3. Functional richness – calculated in R using package *FD* based on traits reflecting pollinator life strategies: parasitism, foraging preference, larval habitat, larval diet, and migration (Table A.5). Species were assigned traits using published sources (Bos et al., 2006; Peeters et al., 2012; Reemer et al., 2009), and functional diversity was computed using a trait-distance framework accounting for species abundances.

The traits included were selected before starting fieldwork and were chosen to represent the different life strategies of the pollinator groups (Table 2). Lastly, pollinator abundance per visit was obtained by adding up the total number of pollinators found with both methods for each visit to a roof.

**Table 2.** Traits. For explanation of traits, see appendix A.5.

| Traits | Categories |
| --- | --- |
| Parasitism | Non-parasitic – Parasitic |
| Foraging preference | Polylectic (no specific forage plant preference) – Oligolectic (specific forage plant preference) |
| Parasitism | Parasitic – non-parasitic |
| Larval habitat | Cavity nesting – Soil nesting – Both cavity & soil nesting – Aquatic free living – Terrestrial free living |
| Larval diet | Arthropod (predators) – Detritus (detritivorous) – Plants (Phytophagous) – Pollen (adults feed larva with pollen) |
| Migration | Non-migratory – migratory |

### Data analysis

To analyse the results, a Bayesian approach to Generalized Linear Models (GLM) was performed using Integrated Nested Laplace Approximation (INLA) with the package INLA in R (Martins et al., 2013; Rue et al., 2009). INLA was used to account for spatial autocorrelation between roofs. Three full models were made, one for each response variable: (1) species richness, (2) functional richness and (3) abundance, with all variables mentioned in Table 1 included as explanatory variables. For pollinator functional richness, a Gaussian model was fitted, while for the count variables pollinator species richness and abundance, a Poisson and a negative binomial model were fitted each respectively. Sampling period, meaning the three periods in which all 25 roofs were visited, was included as a random effect in the model of pollinator abundance to account for the seasonal variation in abundance. Flower seasonality was excluded from the pollinator abundance model, since it was calculated per roof and cannot be calculated per visit. The effect of the weather variables, being temperature, wind speed, sunshine duration and cloud cover, on the response variables was tested beforehand using linear regression. Temperature influenced pollinator abundance and was included in the full model for that response variable. All other weather variables were excluded, because they showed no effect on the response variables. Explanatory variables of all three models were tested for outliers, as well as collinearity, and variables with a Variance Inflation Factor (VIF) higher than three were excluded. Consequently, microrelief, difference in vegetation height, flower diversity, flower species richness and total of vertical structures were dropped in all three models due to high collinearity with each other and type of green roof, as expected. The amount of suitable habitat was correlated with distance to suitable habitat, so only distance was used.

All remaining variables were scaled and included in the full models. Based on these full models, a selection was made based on the DIC. Covariates with the lowest impact were removed one by one until their removal caused the DIC to increase by more than two. Thus, the most parsimonious model within a DIC range of two was selected as the best model. Subsequently, parameters were added and included in additional models if they did not increase DIC with more than two, and were informative, meaning that zero was outside the 95 percent confidence interval. This was done to avoid reporting overly complex models with only one, insignificant additional parameter (Berg et al., 2004). Stochastic Partial Differentiation Equation was used to estimate the spatial autocorrelation in the data and added to the model. All models were unaffected by spatial autocorrelation, as DIC did not improve beyond the range of two. Finally, to see whether species composition differs between types of green roofs, a Venn diagram was made. Nonmetric multidimensional scaling (NMDS) was used to visualize differences in functional traits between the three roof types and a permanova was used to test for significant differences.

Additionally, the analysis was repeated separately for nesting pollinators (all Aculeata excluding Formicidae) and non-nesting pollinators (including Syrphidae, Bombylidae, Conopidae, Stratiomyidae, Rhopalocera, Zygaenidae, and Macroglossinae). Functional richness was not calculated due to limited trait data. The same modelling approach was used for overall pollinator abundance and richness, except a Poisson model was fitted for non-nesting pollinator abundance due to lower dispersion, and the variable ‘bee hotel’ was excluded while vegetation cover was added. These groups were analysed separately to account for their distinct ecological roles and responses to habitat features, allowing for clearer interpretation of group-specific patterns, and interactions with features like bee hotels and vegetation cover.

## Results

### Pollinator community composition

A total of 1199 pollinating insects of 94 species were recorded or collected on the green roofs (Fig. 1). More than half of the individuals were Anthophila bees, including 34 species, and more than a quarter were hoverflies (Syrphidae), including 24 species. A list of all species can be found in the appendix table A.6.

**Figure 1.**
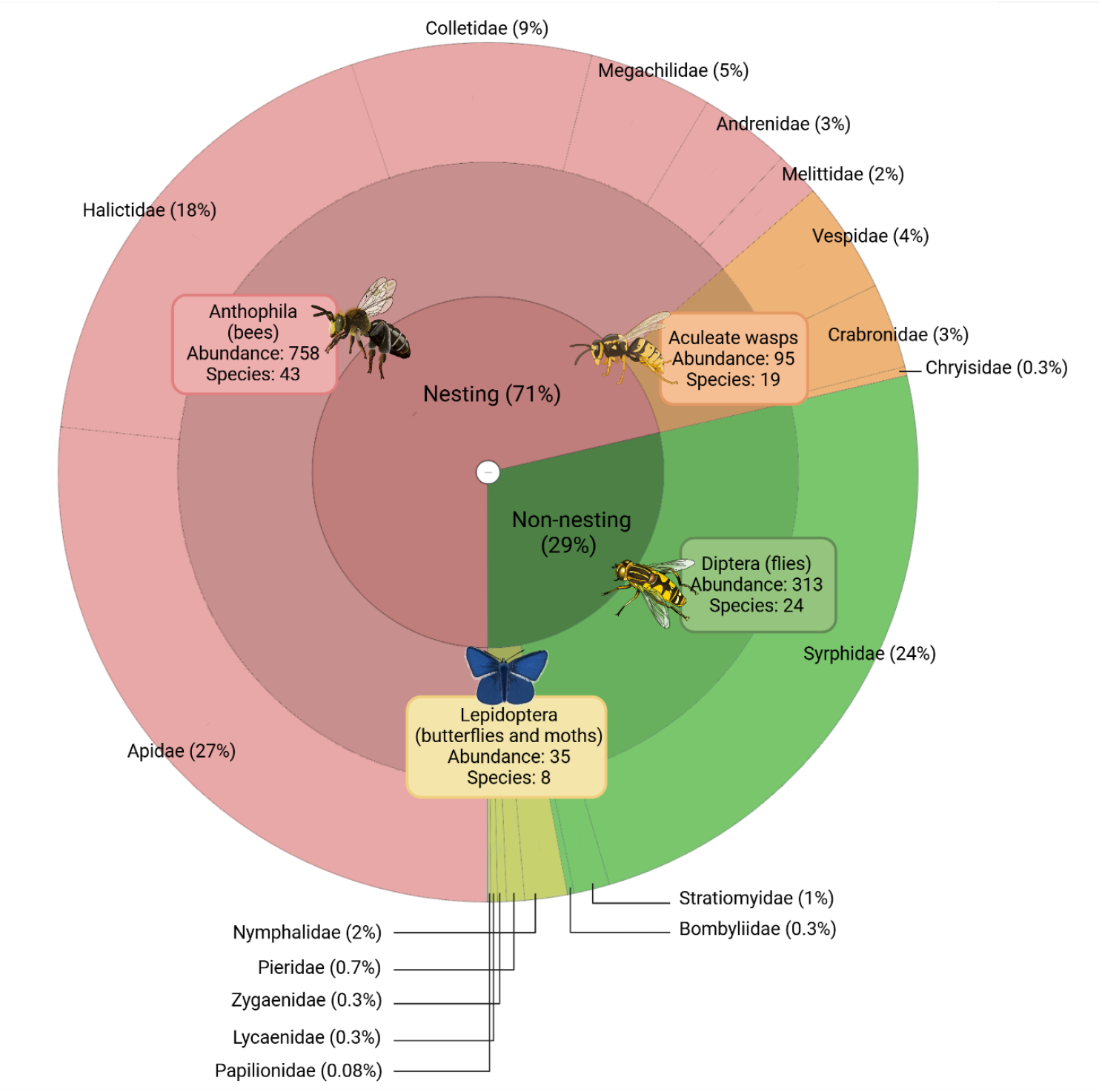
Total abundance (number of individuals) of pollinating insects per family, found on 25 green roofs in urban areas in the Netherlands. The inner circle shows the percentage of nesting vs non-nesting groups. Nesting consists of two groups: Anthophila (bees) and aculeate wasps, of which the abundance and number of species is given in the middle circle. Non-nesting consists of the two groups Diptera (flies) and Lepidoptera (butterflies and day-active moths), also represented with their abundance and number of species in the middle circle. The outer circle shows which families and their abundance in percentages. Pie chart made with Krona and edited with BioRender.com

Six pollinator species considered rare in the Netherlands were found —based on national distribution records (table A.7) (Peeters et al., 2012)— including for example 25 individuals of *Lasioglossum nitidulum* on sedum roofs and 15 of *L. laticeps* on nature roofs and gardens. Eight common pollinator species including five bees and three hoverflies were also recorded (table A8). Common bees included *Hylaeus hyalineatus, L. morio*, and three *Bombus* species. Hoverflies included *Eristalis tenax, Eupeodes corollae*, and *Eupeodes luniger*. These eight common species were found on all roof types.

### Pollinator traits

Most species were common polylectic types, making up 89.5% of species and 97.7% of individuals (Fig. 2). Parasitic species comprised 9% (1.1% in abundance), and 39% had predatory larvae. About 23.4% were migratory, a trait exclusively associated with hoverflies and butterflies in this research.

**Figure 2.**
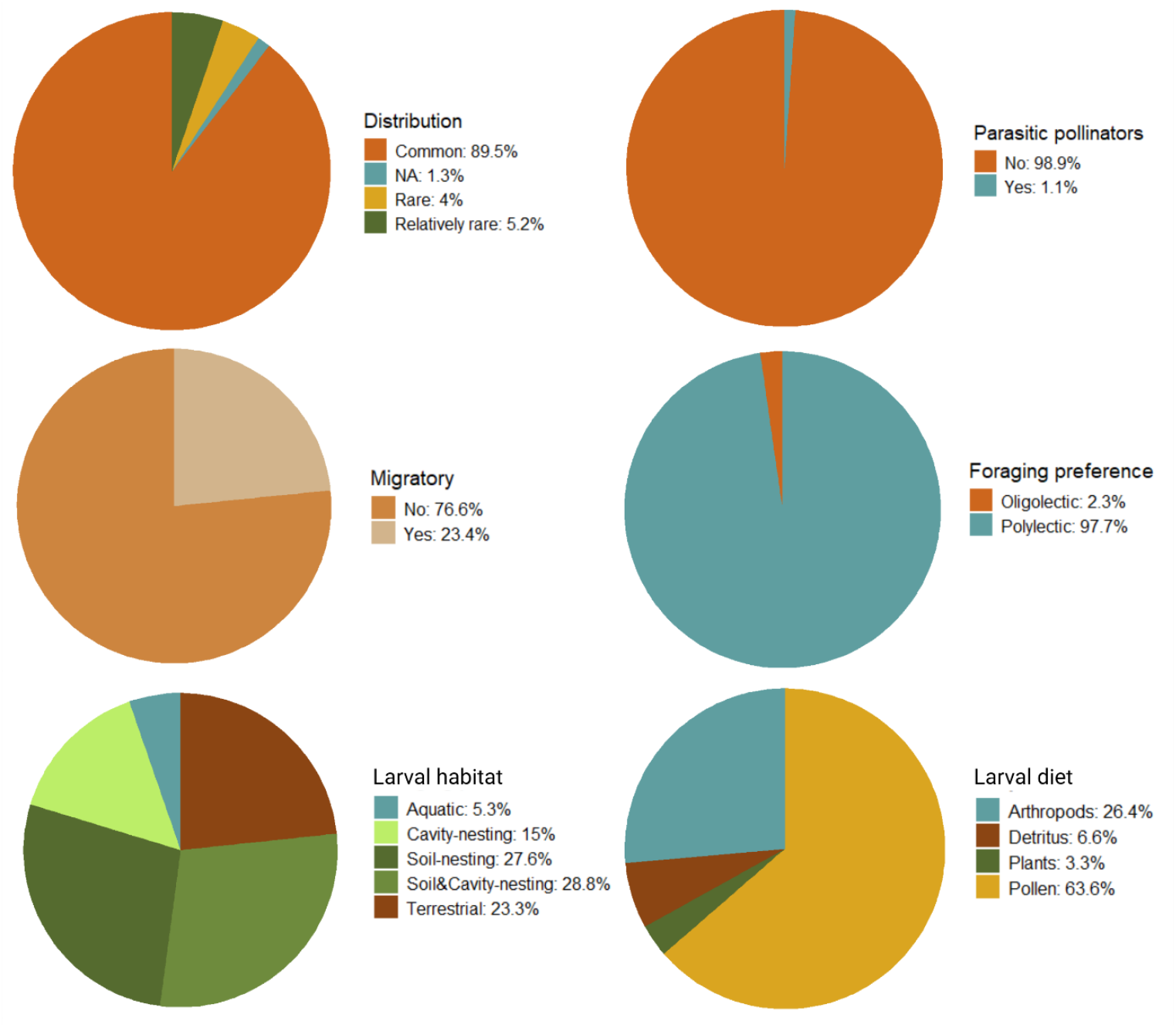
Percentage of abundance of six pollinator traits on green roofs.

Species composition varied by roof type; sedum roofs had the fewest unique species (9), while nature roofs and gardens had 21 and 20, respectively. Functional trait analysis showed no significant difference (PERMANOVA, p = 0.31; NMDS can be found in the appendix fig. B.2).

### Factors influencing pollinator diversity

The generalized linear mixed models identified six key factors that influenced pollinator indices: flower seasonality, flower abundance, height, distance to green space, bee hotels, and honeybee hives. Height and distance were significant only when separating nesting from non-nesting pollinators.

#### (a) Roof type: sedum roofs, nature roofs and roof gardens

Roof type significantly influenced pollinator abundance and functional richness (Fig. 4a and b). Nature roofs significantly supported more than twice as many pollinators as sedum roofs (a 2.16-fold increase; 95% CI: 1.46 to 3.23), and had a significantly higher functional richness (a 1.07-fold increase; 95% CI: 1.02–1.12) compared to sedum roofs. Roof gardens were also found to have significantly higher pollinator abundance (a 2.58-fold increase; 95% CI: 1.81–3.71) and functional richness (a 1.03-fold increase; 95% CI: 0.99–1.08) compared to sedum roofs. These were based on the log-transformed model outputs, which can be found in the appendix table A.9 and A.10.

**Figure 4.**
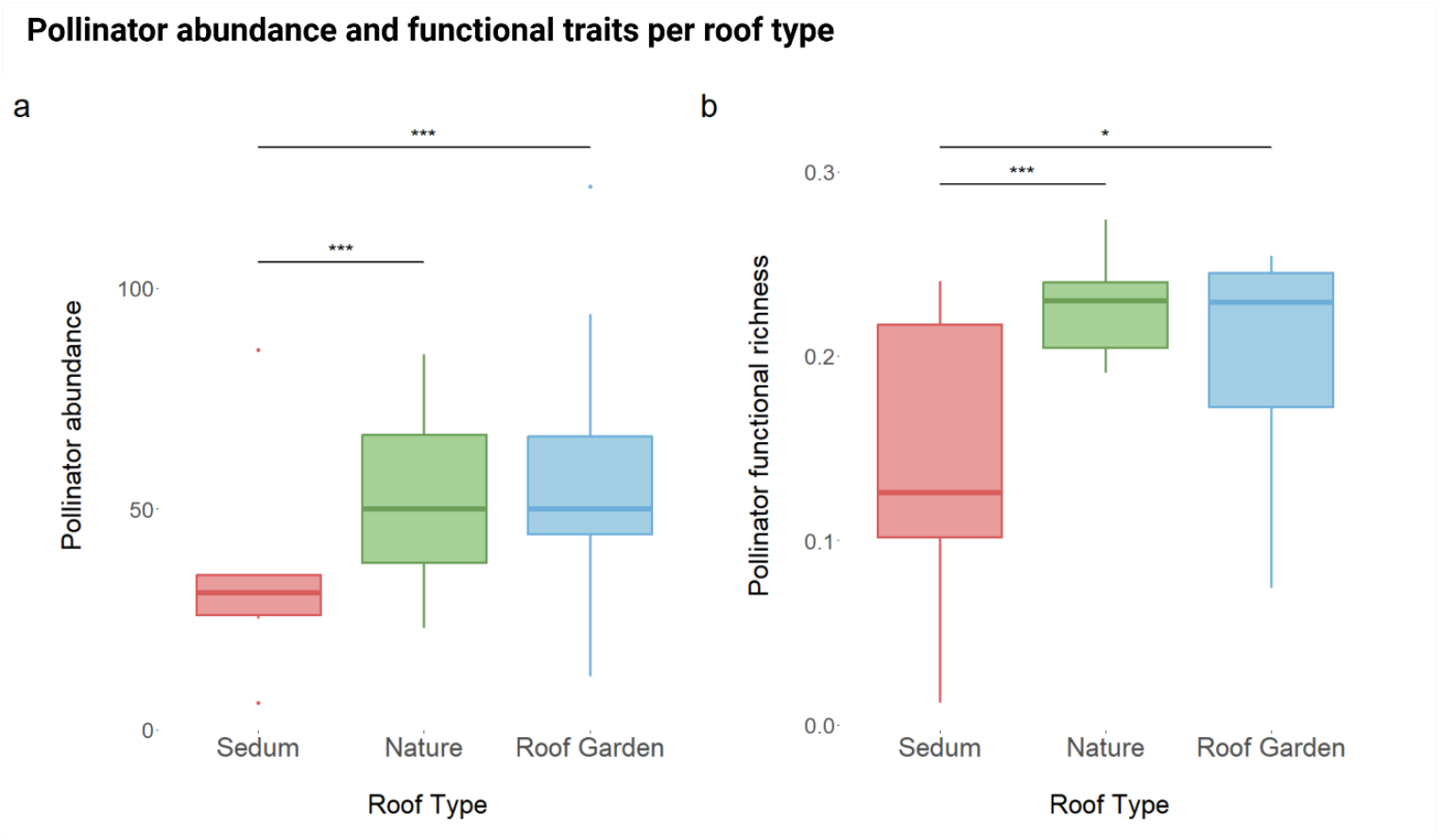
Boxplots of pollinator abundance (a) and functional richness (b) per roof type. Asterisks () indicate significant differences from sedum roofs based on model-estimated contrasts and 95% confidence intervals, ***<0.001 *<0.05. No significant difference was detected between nature roofs and roof gardens. Significance was determined based on the output of the fitted model; no additional post-hoc test was performed.

#### (b) Flower seasonality and abundance

Both flower seasonality and flower abundance were found to have a significant positive effect on pollinator diversity and abundance (Fig. 5).

**Figure 5.**
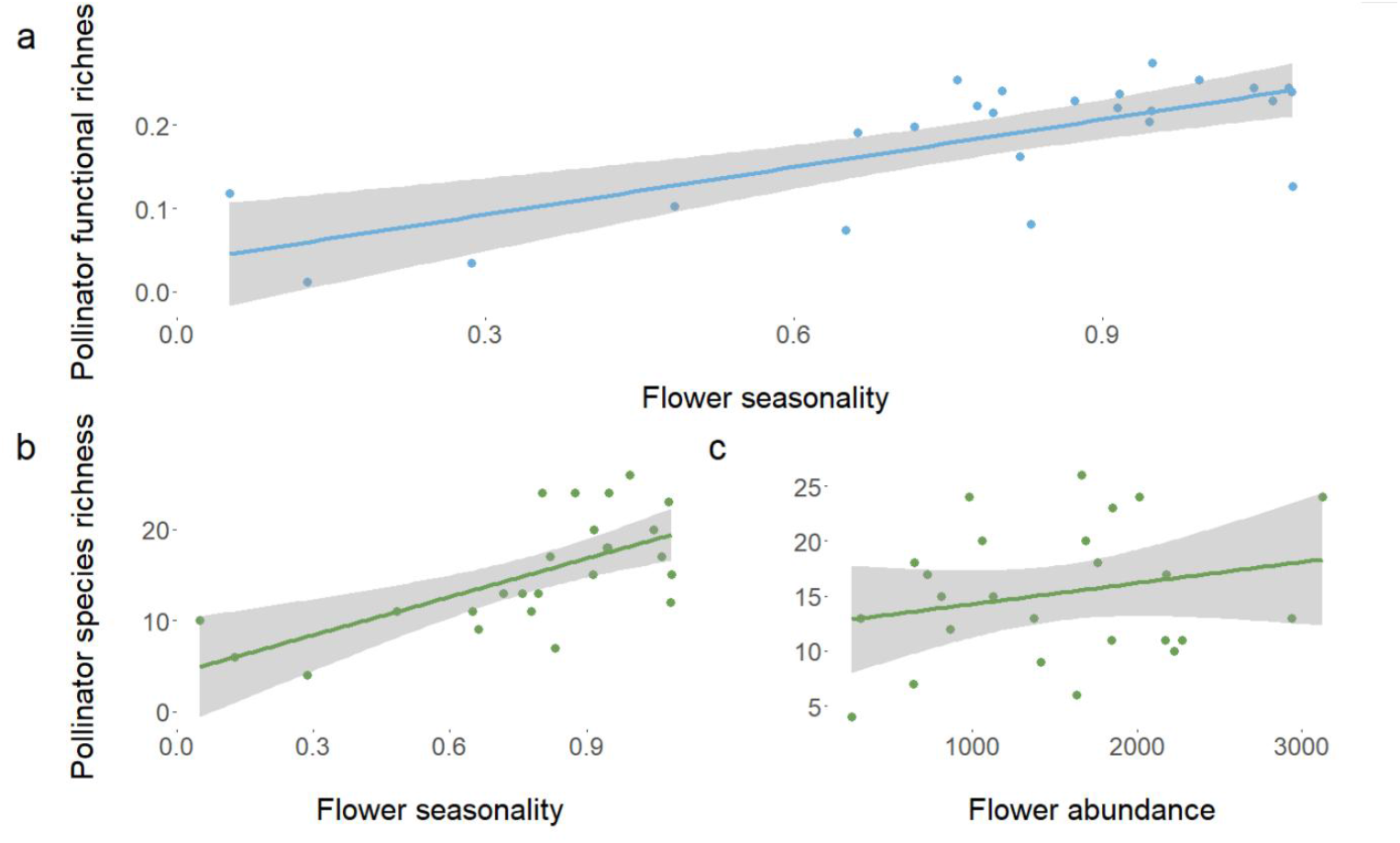
Relationship between (a) Pollinator functional richness and flower seasonality, (b) Pollinator species richness and flower seasonality, (c) Pollinator abundance and flower abundance and (d) Pollinator species richness and flower abundance.

Flower seasonality, defined as a constant supply of flowering plants throughout all three visits (measured by the Shannon index), had a strong positive effect on pollinator species richness (a 3.61-fold increase; 95% CI: 2.25–6.00) and functional richness (a 1.18-fold increase; 95% CI: 1.10–1.27), based on log-linear model estimates. In other words, a more constant supply of flowering plants was associated with a higher pollinator diversity. Additionally, flower abundance positively influenced pollinator species richness (a 1.82-fold increase; 95% CI: 1.21–2.72). Model estimates and confidence intervals can be found in the appendix, table A.9 and A.11. Early spring flower abundance is likely driving the high positive effect of flower seasonality, as many sedum roofs lack flowering plants during this period (Fig. 6).

**Figure 6.**
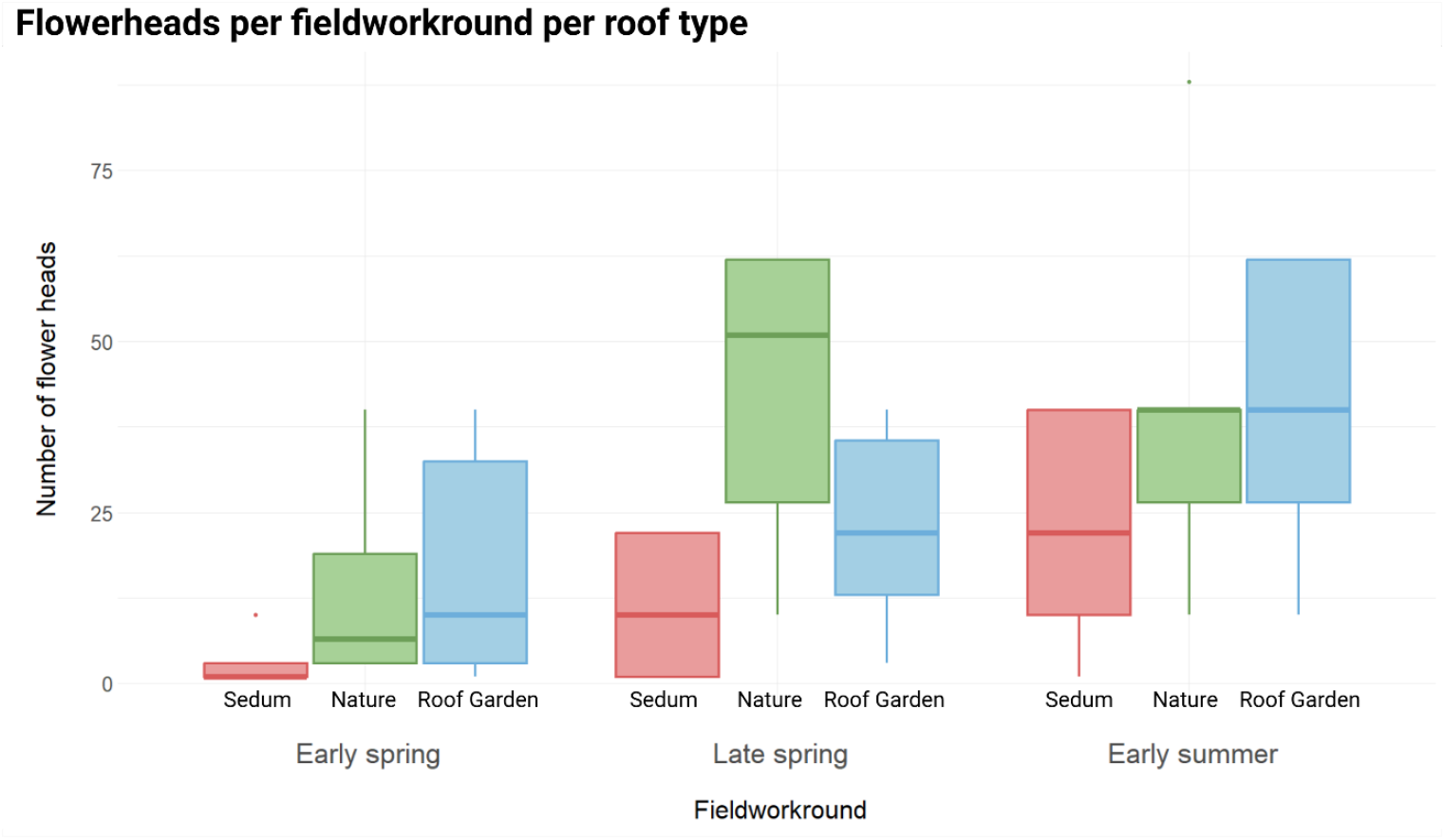
Number of flower heads of all roofs averaged per fieldwork round for each roof type. For each fieldwork round from left to right, it displays Sedum roofs (red), nature roofs (green) and Roof gardens (blue). Early spring = between 8^th^ of April and 15^th^ of May; late spring is between 22^nd^ of May and 4^th^ of June; early summer is between 1^st^ of July and 1^st^ of August.

#### (c) Other factors: honeybee hives, bee hotels, height and distance to surrounding green

These factors were identified as significant by the models (outputs are given in tables A.9 to A.16), but are only briefly discussed here, as certain limitations make the findings less conclusive. Bee hotels and honeybee hives influenced pollinator diversity but were present on only 4 of 25 roofs (see appendix Fig. B.3). Roof height negatively affected nesting but positively affected non-nesting pollinators. (Fig. B.4). However, most of our roofs were between 10 and 25 meters, with two “outliers”; a sedum roof of 40 and a roof garden of 70 meters. Distance from green space was negatively correlated with non-nesting pollinator abundance, though two distant sedum roofs skewed results (>600 m and >1000 m) (Fig. B.5). Due to this, we consider these findings debatable and discuss them further below.

## Discussion

Sedum roofs offer a less suitable habitat for pollinators than nature roofs and roof gardens, likely due to their limited flowering diversity and seasonal gaps in bloom. Although the study provides only a temporal snapshot, it offers insight into the ecological traits and community structure typical of green roof pollinators.

### 1. Pollinator community composition of green roofs

Green roofs support a distinct urban pollinator community, primarily composed of common, generalist polylectic species that collect pollen from a wide range of unrelated plant species. Many of the individuals recorded were species that are often found in urban areas according to the Dutch citizen science platform waarneming.nl, indicating that green roofs form an important connection to the surrounding. This pattern aligns with findings across other urban systems, where cities tend to filter out specialist species, leaving communities dominated by generalist, polylectic taxa (Baldock et al., 2015; Harrison et al., 2018). We observed trophic interactions, such as the presence of *Lasioglossum leucopus* and *L. morio* with their brood parasite *Sphecodes geofrellus* present on the same roof as well as *Cerceris rybyensis*, a specialist predator of *Andrena, Halictus*, and *Lasioglossum* species. These interactions suggest that even relatively small green roofs can support basic ecological complexity and potential interspecific interactions.

The community composition that was found on green roofs in our study confirm previous findings, showing high proportions of polylectic bees (97%) and few parasitic individuals (1.1% abundance, 9% species) (Hofmann & Enner, 2017; Kratschmer et al., 2018). These proportions may be typical for green roofs, where environmental filtering and limited resources favor widespread and adaptable species.. We could indicate key “green roof species” like *Hylaeus hyalinatus* and *L. morio* that are common, consistent with other European green roof studies (Jacobs et al., 2023; Kratschmer et al., 2018; Passaseo et al., 2020), with *L. morio* even found in emerging from the roofs, using emergence traps (Passaseo et al., 2020). Notably, rare or specialist species were also present, such as *Megachile rotundata*, a thermophilic species, and the oligolectic *Macropis europea*, found near its host plant *Lysimachia* on a roof in Amsterdam. This highlights the potential of green roofs to support specialist taxa under the right conditions

### 2. Green roofs as a habitat for reproduction and nesting

Ground-nesting bees were common, indicating that shallow green roof substrates can support nesting. However, sedum roofs seem less suitable (fig. 3) as it had fewer soil-nesters, likely due to their shallower, less complex substrate. This is in line with other studies that found higher percentages of soil-nesting species compared to cavity nesting species (Kratschmer et al., 2018; Ksiazek et al., 2014; Tonietto et al., 2011). It is possible though that ground-nesting bees come from the surrounding and fly up to the rooftops to forage (Tonietto et al., 2011). Yet, for many species the foraging range is not very large with only few species able to cover long distances (Zurbuchen et al., 2010).

It was debated that the high abundance of ground-nesting bees could also be a collecting bias as these species are more likely to be caught with pan traps (Hofmann & Enner, 2017). However, in our study we also used a hand net, so this collecting artifact might not be the case. Cavity-nesters, like *Heriades truncorum*, were more abundant on roofs with bee hotels, highlighting their value in enhancing bee diversity.

Hoverflies were underrepresented on green roofs, with a bee-to-hoverfly ratio of nearly 2:1. This is consistent with findings from other green roof studies, with a 4:1 ratio or even 6:1 ratio (Jacobs et al., 2023; Passaseo et al., 2021). This contrasts with national and ground-level proportions, which are closer to a 1:1 ratio, with some studies reporting a 4:3 (Smit, 2021) or even 3:4 ratio favouring hoverflies (Reemer, 2019). These findings suggest that green roofs are less attractive to hoverflies. But in general urban areas are likely to be less attractive to hoverflies, with lower abundance and species richness of hoverflies compared to bees in urban areas (Baldock et al., 2015). It is suggested that this is due to the homogeneous landscape and lack of suitable habitat to complete their lifecycle (Jacobs et al., 2023; Verboven et al., 2014). Most observed hoverflies were mobile, migratory species, such as *Sphaerophoria scripta*, which may be better able to exploit green roof habitats (Jacobs et al., 2023). Only *S. scripta* has been documented emerging from green roofs (Passaseo et al., 2020), which we found in high numbers (41 individuals) and laying eggs. We did observe one anecdotal sign of reproduction of a hoverfly species; *Paragus haemorrhous* mating and laying eggs. For this hoverfly species, it would be probable that the green roof is used as habitat, as it is a less-mobile species commonly found in dry grasslands (Reemer et al., 2009), which resemble green roofs

### 3. Factors influencing pollinator diversity

#### (a) Roof type

Roof type strongly influenced pollinator richness and abundance. Sedum roofs had significantly lower pollinator diversity compared to nature roofs and roof gardens. This is perhaps unsurprising; sedum roofs, although the most popular green roof option, have limited biodiversity appeal, stemming from their lack of structure and uniform vegetation(Madre et al., 2013; Williams et al., 2014). This would make nature roofs – with only slighter deeper substrates and being cheaper and easier to construct compared to roof gardens – a valuable alternative to sedum roofs.

#### (b) Flower seasonality and abundance

Flower seasonality had a strong positive effect on pollinator species and functional richness. Continuous pollen availability during the flight period is essential (Harris et al., 2024), especially for bees that require nearby forage to provision nests (Zurbuchen et al., 2010). Many sedum roofs lack early-blooming plants (fig. 6), likely driving lower spring pollinator diversity. This can limit bee diversity, especially species active in spring (Matteson et al., 2008) like many Andrena species and bumblebees, which establish colonies in spring (Westphal et al., 2009). As a result, flower scarcity in April and May likely drives lower pollinator richness on these roofs. Indeed, early-season flower abundance was strongly correlated with both species and functional richness, suggesting spring forage availability is a key driver of the positive effect of flower seasonality.

Another thing that stands out, is the high abundance of flowering heads on nature roofs compared to roof gardens in late spring (fig. 6). This makes sense as nature roofs are often dominated by a diverse mix of native and semi-natural plant species adapted to early-season flowering (Sutton et al., 2015). These plants, such as early-flowering herbs and grasses, are typically selected for their ecological value and resilience under low-input conditions, which has been found to positively influence insect diversity (Fenoglio et al., 2023). By contrast, roof gardens—especially those with non-native plants —may include species that have later bloom times (fig.6). Even though non-native plants are often found to be less beneficial to pollinators, several studies find that pollinators also often do not discriminate between native and non-native species (Wenzel et al., 2020). This may be particularly relevant in urban habitats, where pollinator communities appear more flexible: in urban sites, native and non-native plants received similar visitation from pollinators, while in farmland and nature reserves, visits were almost exclusively to native plants (Baldock et al., 2015). Because native and non-native plants tend to bloom at different times, they can complement each other by extending the availability of floral resources across the season—suggesting that even non-native plantings can enhance the ecological value of urban green roofs when thoughtfully integrated.

#### (c) Height and distance to suitable habitat on nesting and non-nesting pollinators

Height negatively affected nesting pollinators, which is consistent with previous studies showing reduced bee diversity on taller buildings (MacIvor, 2016; Tonietto et al., 2011). It is likely driven by the limited mobility of smaller species (MacIvor, 2016; Zurbuchen et al., 2010). In contrast, hoverflies and butterflies—more mobile species—were positively associated with height. However, this may be confounded by roof type. Our highest roof was a roof garden, that generally showed higher diversity in our study — which may better explain the positive association.

There are other green roof studies that found a positive trend of height with insect diversity as well. Kyrö et al. (2020) studied roofs of up to 11 meters and showed that arthropod diversity in general went up. In another study, roofs of up to 56 meters were investigated, showing that…(Lin & Chen, 2022). They hypothesized that reduced competition, presence of highly mobile species recorded, or other roof factors that were accompanied by increasing heights could explain the positive trend. We emphasize that our findings rather suggest that mobile species may not be constrained by heights of up to 70 meters, provided that a resource-rich habitat, such as a roof garden, is available. The trends found here need further investigation, using more replicates of roofs of specific heights.

Distance to green space negatively affected non-nesting pollinator abundance,. Surrounding green is often said to positively influence diversity on green roofs (Dromgold et al., 2020; Hussain et al., 2023; Ksiazek-Mikenas et al., 2018; MacIvor, 2016; Tonietto et al., 2011), so this would align with our findings. But in our study the trend was also not very strong (Estimate -0.719), and more likely driven by two sedum roof outliers (App. Fig. B5), so we do not want to conclude this as a definitive relationship..,

#### (d) Honeybee hives and bee hotels

The negative effects of honeybee hives on pollinator abundance align with existing research, indicating that honeybees can outcompete native pollinators for floral resources (Geldmann & González-varo, 2018; Macinnis et al., 2023; Mallinger et al., 2017; Ropars et al., 2019). This pressure is likely amplified on green roofs due to their isolation and limited forage. Native bees nesting on roofs are especially vulnerable, as they rely heavily on local flowers (Zurbuchen et al., 2010). In contrast, bee hotels may benefit certain species—such as the cavity-nesting *Heriades truncorum*—and could be a more supportive measure for enhancing wild bee species presence. However, due to the limited number of roofs with honeybee hives and bee hotels, the observed effects remain inconclusive and should be further looked into.

## Conclusions

In summary, these findings highlight the importance of constant forage availability, especially in spring. Sedum roofs are a less suitable habitat for pollinators compared to other roof types, which can be attributed to the lack of flowering plants in early spring, as well as the lower diversity of plant species throughout the year. If the goal is to help biodiversity, specifically insects, nature roofs perform better than sedum roofs and would therefore be more useful to implement. While roof height may limit bee presence, it appears less restrictive for butterflies and hoverflies, though green roofs generally attract fewer hoverflies overall. Despite these limitations, Dutch green roofs support a diverse pollinator community, including rare species, underscoring their ecological value in urban environments. Enhancing this value will require prioritizing nature roofs and roof gardens, avoiding new honeybee hive placements, and ensuring diverse, seasonally distributed plantings to maintain forage throughout the year.

## Acknowledgements

We would like to thank our supervisor at Bureau Stadsnatuur, Rens de Boer, help setting up this research, providing with necessary materials and his continuous help and feedback during the entire process. Secondly, we would like to thank Frank van der Meer for helping with the identification of many difficult bee species. Furthermore, we would like to thank Marco Roos for his feedback. Additionally, we would like to thank the ecologists at Bureau Stadsnatuur for their contributions and their presence during identification. In particular, a special thanks to Remko Anderweg for help with plant identification, Sander Elzerman for his feedback on ideas and hypothesises, and Wouter Moerland for assistance with hoverfly identification and substantive discussions on hoverfly and bee ecology and its connection with green roof design. We would like to thank Sytske de Waart for the help with sorting of catches and finding some bees missed when sorting. Moreover, we would like to thank Wim Klein for identifying several wasp species and Bram Langeveld for his help with getting collection material. Finally, many thanks to all the people that let us do research on their green roofs.

