## Supplemental material for "Scarcity of flowering plants on *Sedum* roofs limits pollinator diversity"

### Supplementary material for the article: Scarcity of flowering plants on *Sedum* roofs limits pollinator diversity

#### Appendices A (for tables) and B (for figures)

##### A. Supplementary Tables

Table A.1. Species groups

| Group | Common name |
|---|---|
| Apoidea: Apidae | Bees |
| Apoidea: excl. Apidae | Digger wasps |
| Chrysidinae | Cuckoo wasps |
| Vespidae (excl. Formicidae) | Wasps (Dutch: Plooivleugelwespen) |
| Rhopalocera | Butterflies |
| Macroglossinae | Day-active hawk-moths |
| Zygaenidae | Burnet moths |
| Syrphidae | Hoverflies |
| Bombylidae | Bee flies |
| Conopidae | Thick-headed flies |
| Stratiomyidae | Soldier flies |

Table A.2. Green roof locations with coordinates and type of roof

| **Roof** | **Municipality** | **Type of roof** | **Intensive - extensive** | **X-coordinate** | **Y-coordinate** |
|---|---|---|---|---|---|
| Accenture | Amsterdam | Roof garden | Intensive | 52.33735 | 4.87299 |
| Aegon | Den Haag | Roof garden | Intensive | 52.09223 | 4.37036 |
| B.Amsterdam | Amsterdam | Roof garden | Intensive | 52.34314 | 4.82878 |
| CBK | Rotterdam | Sedum | Extensive | 51.91560 | 4.47614 |

| | | | | | |
|---|---|---|---|---|---|
| Dakakker | Rotterdam | Roof garden | Intensive | 51.92537 | 4.47672 |
| Dalton | Leidschendam-Voorburg | Sedum | Extensive | 52.07608 | 4.37175 |
| De Boele | Amsterdam | Roof garden | Intensive | 52.33474 | 4.87414 |
| Eneco | Rotterdam | Roof garden | Intensive | 51.95355 | 4.55781 |
| Erasmus MC | Rotterdam | Roof garden | Intensive | 51.91118 | 4.46718 |
| Gemeentehuis | Hellevoetssluis | Sedum | Extensive | 51.82384 | 4.13037 |
| Het Nieuwe Instituut | Rotterdam | Sedum | Extensive | 51.91489 | 4.47069 |
| IVN Hortus | Amsterdam | Sedum | Extensive | 52.36711 | 4.90834 |
| Lijnbaanflat Joost Banckertsplaats | Rotterdam | Sedum | Extensive | 51.92143 | 4.47503 |
| Kalverpassage | Amsterdam | Sedum | Extensive | 52.36751 | 4.89160 |
| Lumen | Wageningen | Sedum | Extensive | 51.98776 | 5.66764 |
| Museon | Den Haag | Nature | Extensive | 52.08862 | 4.28074 |
| NEMO | Amsterdam | Nature | Extensive | 52.37452 | 4.91229 |
| NIOO | Wageningen | Nature | Extensive | 51.98715 | 5.67119 |
| Nieuwe Universiteitsgebouw VU | Amsterdam | Nature | Extensive | 52.33437 | 4.86369 |
| Smartroof 2.0 | Amsterdam | Nature | Extensive | 52.37224 | 4.91569 |
| Stadskantoor | Helmond | Nature | Extensive | 51.47631 | 5.65802 |
| Strijp-S 'Anton' | Eindhoven | Roof garden | Intensive | 51.44765 | 5.45538 |
| Theil Building | Rotterdam | Sedum | Extensive | 51.91878 | 4.52578 |
| VU Dakterras | Amsterdam | Roof garden | Intensive | 52.33409 | 4.86612 |
| Groot Willemsplein | Rotterdam | Roof garden | Intensive | 51.91055 | 4.48139 |

Table A.3. Guidelines for counting inflorescences, as an indicator for flower abundance.

| Plant | Inflorescence type | Explanation* |
|---|---|---|
| Asteraceae | Heads | |
| Eupatorium | Umbel | |

| | | |
|---|---|---|
| Caprifoliaceae | Heads | |
| Trifolium & Medicago | Head | Globose heads count as one unit |
| Lotus | Umbel | Umbels with 2-6 flowers arranged in a circle count as one unit |
| Apiaceae | Umbel | |
| Boraginaceae | Scorpioid cyme | Coiled cyme, uncoils as it flowers: whole cyme, both coiled an uncoiled part, count as one unit |
| Brassicaceae | Flower stalk | Whole flower raceme count as one unit |
| Onograceae | Flower stalk | Whole flower raceme count as one unit |
| Orobranchaceae | Flower stalk | Whole stalk count as one unit |
| Lamiaceae | Flower stalk | Whole stalk count as one unit |
| Orchidaceae | Flower stalk | Whole stalk count as one unit |
| Polygonaceae | Spike | |
| Plantago spec. | Spike | |
| Veronica spec | Single flower | |
| Geraniaceae | Single flower | |
| Non-woody Rosaceae | Single flower | |
| Woody Rosaceae | Single flower | Estimate number of flowers per branch, than multiply with number of branches |
| Filipendula spec. | Corymb | Whole compound corymb count as one unit |
| Caryophyllaceae | Single flower | |
| Ranunculaceae | Single flower | |
| Lysimachia spec. | Compound raceme | Whole terminal compound raceme count as one unit |
| Primulaceae | Single flower | |
| Rubiaceae | Cyme | Whole cyme count as one unit |
| Buddleja spec. | Panicle | |
| Other | nvt | Count what is practical |

Table A.4: Weather variables

| Group of variable | Variables | Unit | Method |
|---|---|---|---|
| Weather | Temperature | Celsius | From KNMI |
| | Sunshine duration (during pan trapping) | Hours | From KNMI |
| | Cloud cover (during catching) | In parts of one/eight | Estimated in the field |
| | Wind speed | Km/u | From KNMI |

Table A.5: Explanation of traits

| Traits | Categories | Explanation | Example genus |
|---|---|---|---|
| Parasitism | Non-parasitic | Larvae forage independently or are provisioned by parents | Andrena |
| | Parasitic | Larvae eat provisioning and/or larvae of other species | Nomada |
| Foraging preference | Polylectic | Imago’s feed of a wide variety of plants for pollen and nectar, show no clear preference | Eristalis |
| | Oligolectic & monolectic | Imago’s exhibit clear preference for one genus or species of plant to collect pollen | Macropis |
| Larval habitat | Soil-nesting | Makes nest in the ground | Halictus |
| | Cavity-nesting | Makes nest in other structures above the ground | Megachile |
| | Both soil & cavity nesting | Can make nests both above and below the ground | Many species of Bombus |
| | Aquatic | Larvae live in water | Helophilus |
| | Terrestrial | Larvae live on land | Platycheirus |
| Food larvae | Pollen | Feed on pollen | Osmia |
| | Predatory | Feed on animals | Ancistrocerus |
| | Phytophagous | Feed on plants | Eumerus |

| | | | |
|---|---|---|---|
| | Saprophagous | Feed on rotting, dead or waste material | Rhingia |
| Migration | Non-migratory | Does not migrate | Paragus |
| | Migratory | Migrates | Vanessa |

Table A.6: Species list

| **Anthophila (bees)** | **Syrphidae (hoverflies)** | **Aculeata (excl. Anthophila) (wasps)** | **Other** |
|---|---|---|---|
| Andrena barbilabris | Cheilosia latifrons | Ancistrocerus parietum | Aricia agestis |
| Andrena bicolor | Episyrphus balteatus | Cerceris rybyensis | Celestrina argiolus |
| Andrena chrysosceles | Eristalis arbustorum | Crabronidae spec. | Chloromyia formosa |
| Andrena flavipes | Eristalis intricaria | Crossocerus elongatus | Papillio machaon |
| Andrena fulva | Eristalis nemorum | Crossocerus ovalis | Pieris brassicae |
| Andrena labiata | Eristalis tenax | Crossocerus wesmaili | Pieris rapae |
| Andrena minitula | Eupeodes corollae | Hedychrum niemelai | Vanessa atalanta |
| Andrena mitis | Eupeodes luniger | Lindenius albilabris | Vanessa cardui |
| Andrena nigroaenea | Helophilus pendulus | Mimumesa ovalis | Villa hottentotta |
| Andrena nitida | Helophilus trivitattus | Mimumesa unicolor | Zygaena filipendulae |
| Bombus hortorum | Melanostoma mellinum | Oxybelus bipunctata | |
| Bombus hypnorum | Merodon equestris | Passaloecus pictus | |
| Bombus lapidarius | Paragus haemorrhous | Phlanthus triangulum | |
| Bombus pascuorum | Pipizella viduata | Polistes dominulus | |
| Bombus pratorum | Platycheirus albimanus | Pseudomalus auratus | |
| Bombus terrestris-complex | Platycheirus scutatus-complex | Pseudomalus violaceus | |
| Bombus lucorum | Scaeva pyrastri | Trypoxylon figulus | |
| Chelostoma rapunculi | Spaerophoria ruepelli | Vespula germanica | |
| Colletes daviesanus | Sphaerophoria scripta | Vespula vulgaris | |
| Dasypoda hirtipes | Syrphus ribesii | | |

| | |
|---|---|
| Halictus tumulorum | Syrphus torvus |
| Heriades tncorum | Syrphus vitripennis |
| Hylaeus communis | |
| Hylaeus hyalineatus | |
| Hylaeus pictipes | |
| Lasioglossum calceatum | |
| Lasioglossum laticeps | |
| Lasioglossum leucopus | |
| Lasioglossum leucozonium | |
| Lasioglossum minitissimum | |
| Lasioglossum morio | |
| Lasioglossum nitidulum | |
| Lasioglossum sexstrigatum | |
| Macropis europaea | |
| Megachile rotundata | |
| Megachile willughbiella | |
| Megachile centuncularis | |
| Nomada fucata | |
| Nomada goodeniana | |
| Nomada sheppardana | |
| Osmia bicornis | |

| |
|---|
| Sphecodes geofrellus |
| Sphecodes miniatus |

Table A.7. Rare species, with the number of roofs they were found on and in brackets the abundance per roof in percentage.

| Species | Family | Number of observations | Number of roofs (species abundance%) per roof type | | |
|---|---|---|---|---|---|
| | | | Sedum | Biodiverse | Roof Garden |
| *Lasioglossum laticeps* | Halictidae | 15 | 1 (7%) | 1 (40%) | 1 (53%) |
| *Lasioglossum nitidulum* | Halictidae | 25 | 3 (92%) | | 1 (8%) |
| *Megachile rotundata* | Megachilidae | 2 | | 1 (100%) | |
| *Sphecodes geoffrellus* | Halictidae | 2 | | | 2 (100%) |
| *Hylaeus pictipes* | Colletidae | 1 | 1 (100%) | | |
| *Passaloecus pictus* | Crabronidae | 2 | | 2 (100%) | |

Table A.8. Common species (meaning > 50 observations), the number of roofs they were found on and in brackets the abundance per roof in percentage.

| Species | Family | Number of observations | Number of roofs (species abundance%) per roof type | | |
|---|---|---|---|---|---|
| | | | Sedum | Biodiverse | Roof Garden |
| *Hylaeus hyalineatus* | Halictidae | 98 | 8 (58%) | 3 (22%) | 8 (20%) |
| *Lasioglossum morio* | Halictidae | 99 | 6 (20%) | 5 (42%) | 6 (38%) |
| *Bombus lapidarius* | Apidae | 103 | 3 (23%) | 4 (28%) | 10 (49%) |
| *Bombus pascuorum* | Apidae | 68 | 6 (25%) | 5 (26%) | 8 (49%) |
| *Bombus terrestris-complex* | Apidae | 127 | 7 (22%) | 6 (29%) | 10 (49%) |
| *Eristalis tenax* | Syrphidae | 57 | 3 (10%) | 4 (25%) | 8 (65%) |
| *Eupeodes corollae* | Syrphidae | 54 | 6 (37%) | 4 (15%) | 7 (48%) |
| *Eupeodes luniger* | Syrphidae | 51 | 5 (18%) | 3 (23%) | 9 (59%) |

Table A.9: Selected models for **pollinator functional richness** using INLA

| | **Model 1** | | | **Model 2** | | |
|---|---|---|---|---|---|---|
| | Estimate | 2.5% | 97.5% | Estimate | 2.5% | 97.5% |
| Flower seasonality | 0.166 | 0.095 | 0.237 | 0.146 | 0.074 | 0.218 |
| Honeybee hive | -0.102 | -0.172 | -0.032 | -0.117 | -0.187 | -0.048 |
| Type of roof: nature | 0.070 | 0.023 | 0.117 | 0.090 | 0.039 | 0.141 |
| Type of roof: roof garden | 0.033 | -0.013 | 0.079 | 0.050 | 0.002 | 0.099 |
| Roof age | X | | | 0.065 | -0.012 | 0.141 |
| DIC | 79.6 | | | 80.8 | | |

Table A.10: Selected models for **pollinator abundance** using INLA

| | **Model 1** | **Model 2** |
|---|---|---|

| | Estimate | 2.5% | 97.5% | Estimate | 2.5% | 97.5% |
|---|---|---|---|---|---|---|
| Bee hotel: Yes | 0.411 | 0.026 | 0.815 | 0.429 | 0.042 | 0.838 |
| Honeybee hive: Yes | -1.622 | -2.258 | -0.981 | -1.578 | -2.215 | -0.935 |
| Type of roof: nature | 0.770 | 0.380 | 1.172 | 0.721 | 0.324 | 1.129 |
| Type of roof: roof garden | 0.949 | 0.593 | 1.311 | 0.915 | 0.554 | 1.278 |
| Flower abundance | X | | | 0.548 | -0.123 | 1.266 |
| DIC | 518.6 | | | 518.9 | | |

Table A.11: Selected models for **pollinator species richness** using INLA

| | **Model 1** | | | **Model 2** | | |
|---|---|---|---|---|---|---|
| | Estimate | 2.5% | 97.5% | Estimate | 2.5% | 97.5% |
| Flower abundance | 0.645 | 0.206 | 1.048 | 0.596 | 0.190 | 1.001 |
| Flower seasonality | 1.366 | 0.886 | 1.874 | 1.285 | 0.809 | 1.792 |
| Honeybee hives | X | | | -0.356 | -0.847 | 0.098 |
| DIC | 142.2 | | | 141.8 | | |

Table A.12: Selected models for **species richness of nesting pollinators** using INLA

| | **Model 1** | | | **Model 2** | | |
|---|---|---|---|---|---|---|
| | Estimate | 2.50% | 97.50% | Estimate | 2.50% | 97.50% |
| Flower seasonality | 1.081 | 0.495 | 1.709 | 1.047 | 0.484 | 1.654 |
| Flower abundance | 0.833 | 0.354 | 1.309 | 0.732 | 0.228 | 1.234 |
| Height of the building | -1.142 | -2.125 | -0.185 | X | | |
| Honeybee hive | X | | | -0.615 | -1.393 | 0.07 |
| DIC | 132 | | | 134 | | |

Table A.13: Selected models for **species richness of non-nesting pollinators** using INLA

| | **Model 1** | | | **Model 2** | | |
|---|---|---|---|---|---|---|
| | Estimate | 2.50% | 97.50% | Estimate | 2.50% | 97.50% |
| Flower seasonality | 1.89 | 1.039 | 1.709 | 1.047 | 0.484 | 1.654 |
| Height of the building | 1.171 | 0.444 | 1.309 | 0.732 | 0.228 | 1.234 |

| | | | | | | |
|---|---|---|---|---|---|---|
| Type of roof: roof garden | X | 0.274 | -0.185 | X | | |
| DIC | 103.9 | | | 104.4 | -1.393 | 0.07 |
| Flower seasonality | 1.89 | | | 1.039 | | |

Table A.14: Selected models for **abundance of nesting pollinators** using INLA

| | **Model 1** | | | **Model 2** | | |
|---|---|---|---|---|---|---|
| | Estimate | 2.50% | 97.50% | Estimate | 2.50% | 97.50% |
| Honeybee hive | -1.518 | -2.257 | -0.768 | -1.493 | -2.241 | -0.734 |
| Type of roof: roof garden | 0.945 | 0.518 | 1.379 | 0.849 | 0.435 | 1.269 |
| Type of roof: nature | 0.686 | 0.221 | 1.167 | 0.549 | 0.14 | 1.062 |
| Bee hotel: yes | 0.367 | -0.095 | 0.856 | X | | |
| DIC | 479.9 | | | 480.3 | | |

Table A.15: Selected models for **pollinator functional richness** using INLA

| | Model 1 | | | Model 2 | | |
|---|---|---|---|---|---|---|
| | Estimate | 2.5% | 97.5% | Estimate | 2.5% | 97.5% |
| Flower seasonality | 0.166 | 0.095 | 0.237 | 0.146 | 0.074 | 0.218 |
| Honeybee hive | -0.102 | -0.172 | -0.032 | -0.117 | -0.187 | -0.048 |
| Type of roof: Nature | 0.070 | 0.023 | 0.117 | 0.090 | 0.039 | 0.141 |
| Type of roof: roof garden | 0.033 | -0.013 | 0.079 | 0.050 | 0.002 | 0.099 |
| Roof age | X | | | 0.065 | -0.012 | 0.141 |
| DIC | 79.6 | | | 80.8 | | |

Table A.16. Selected models for **abundance of non-nesting pollinators** using INLA

| | **Model 1** | | | **Model 2** | | |
|---|---|---|---|---|---|---|
| | Estimate | 2.50% | 97.50% | Estimate | 2.50% | 97.50% |
| Height of building | 1.259 | 0.687 | 1.83 | 1.144 | 0.548 | 1.735 |
| Honeybee hive | -0.858 | -1.394 | -0.359 | -0.692 | -1.254 | -0.169 |
| Distance to habitat | -0.719 | -1.427 | -0.047 | -0.735 | -1.446 | -0.061 |

| | | | | | | |
|---|---|---|---|---|---|---|
| Type of roof: roof garden | 0.992 | 0.683 | 1.311 | 0.998 | 0.689 | 1.318 |
| Type of roof: nature | 0.691 | 0.344 | 1.044 | 0.679 | 0.332 | 1.032 |
| Roof area | X | | | 0.284 | -0.055 | 0.62 |
| DIC | 404.8 | | | 404.2 | | |

#### B. Supplementary figures

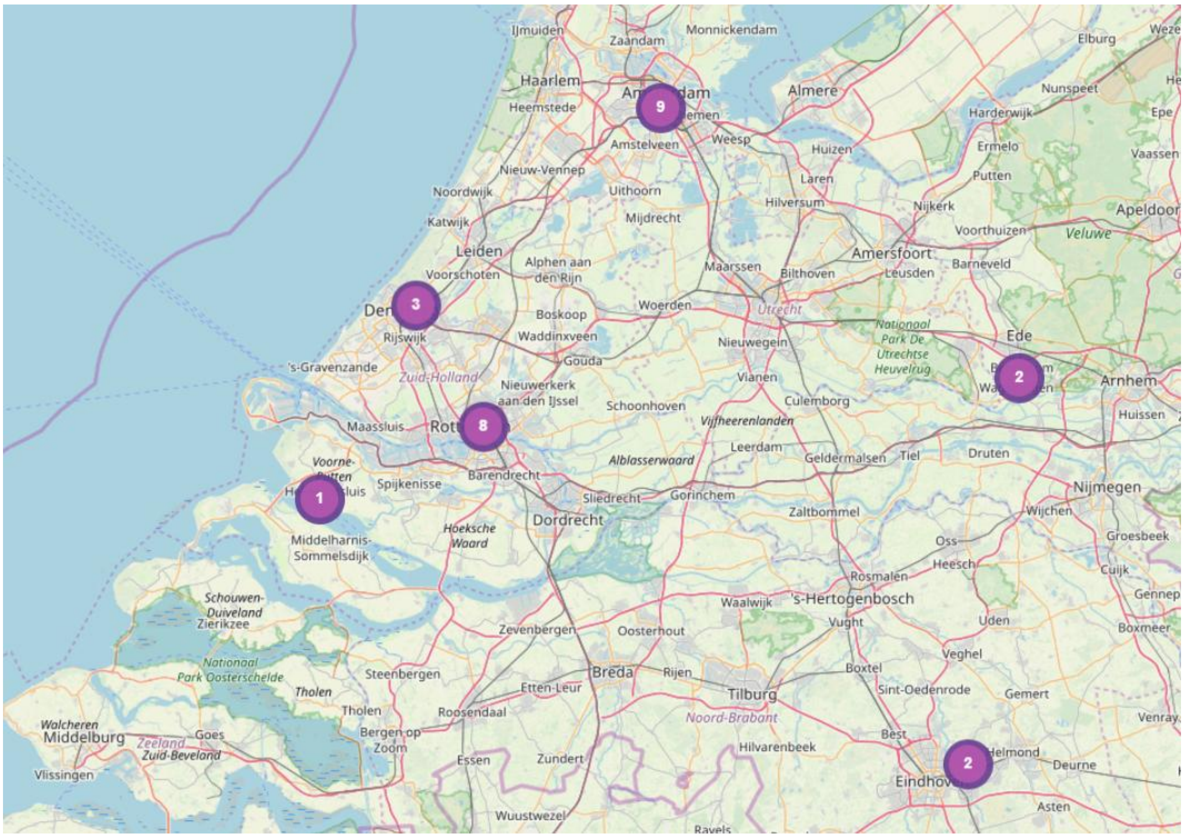

Fig. B.1: Locations of green roofs sampled on map. The number of roofs within a given city is given by the numbers within the circles.

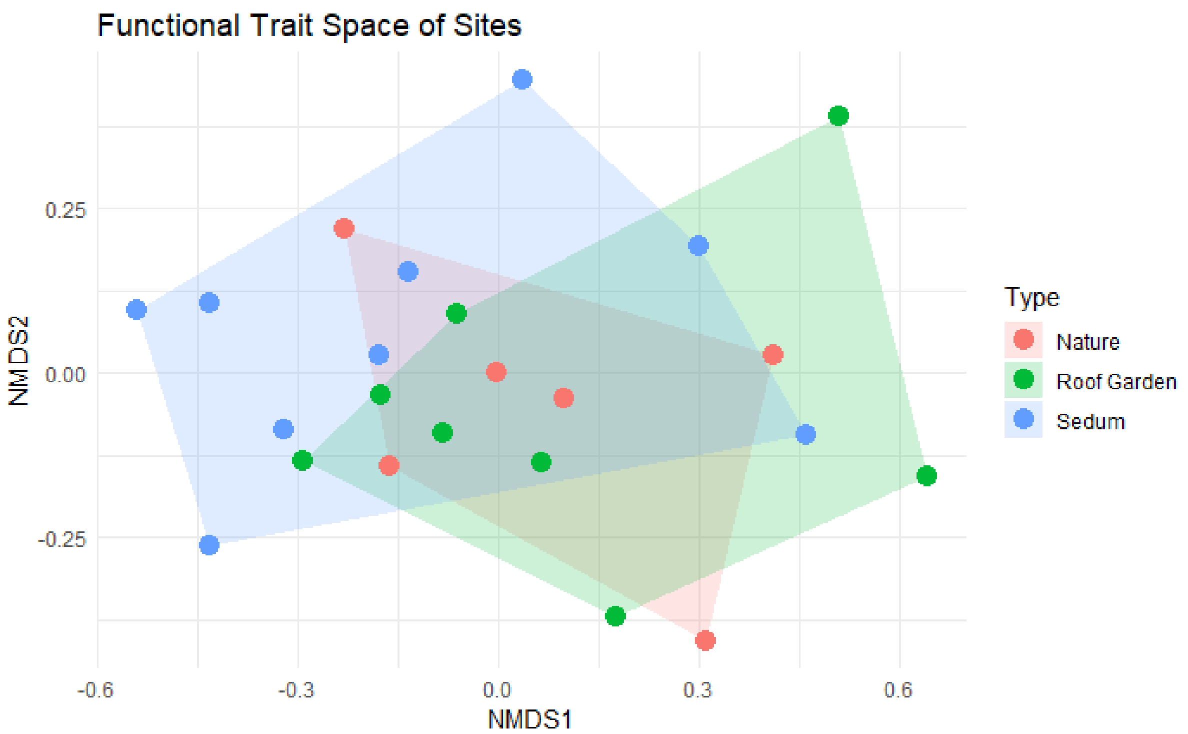

Fig. B.2. NMDS based on functional traits to compare nature roofs, roof gardens and sedum roofs.

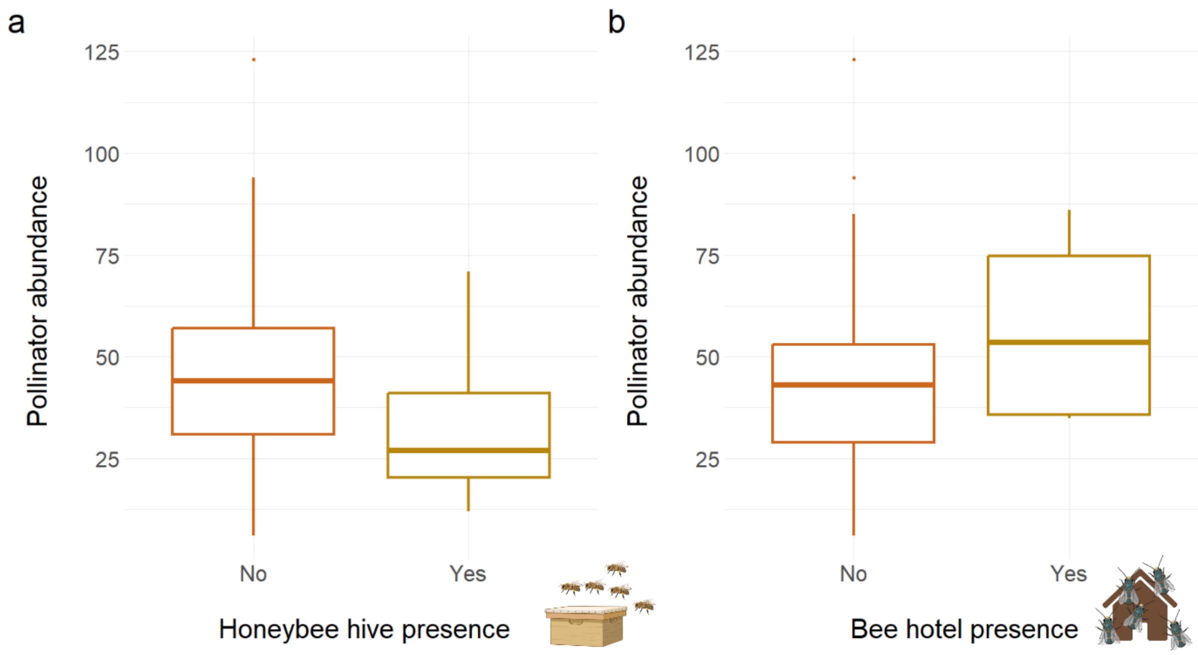

Fig. B.3. Boxplots for pollinator abundance in the presence of (a) honeybee hives and (b) bee hotels

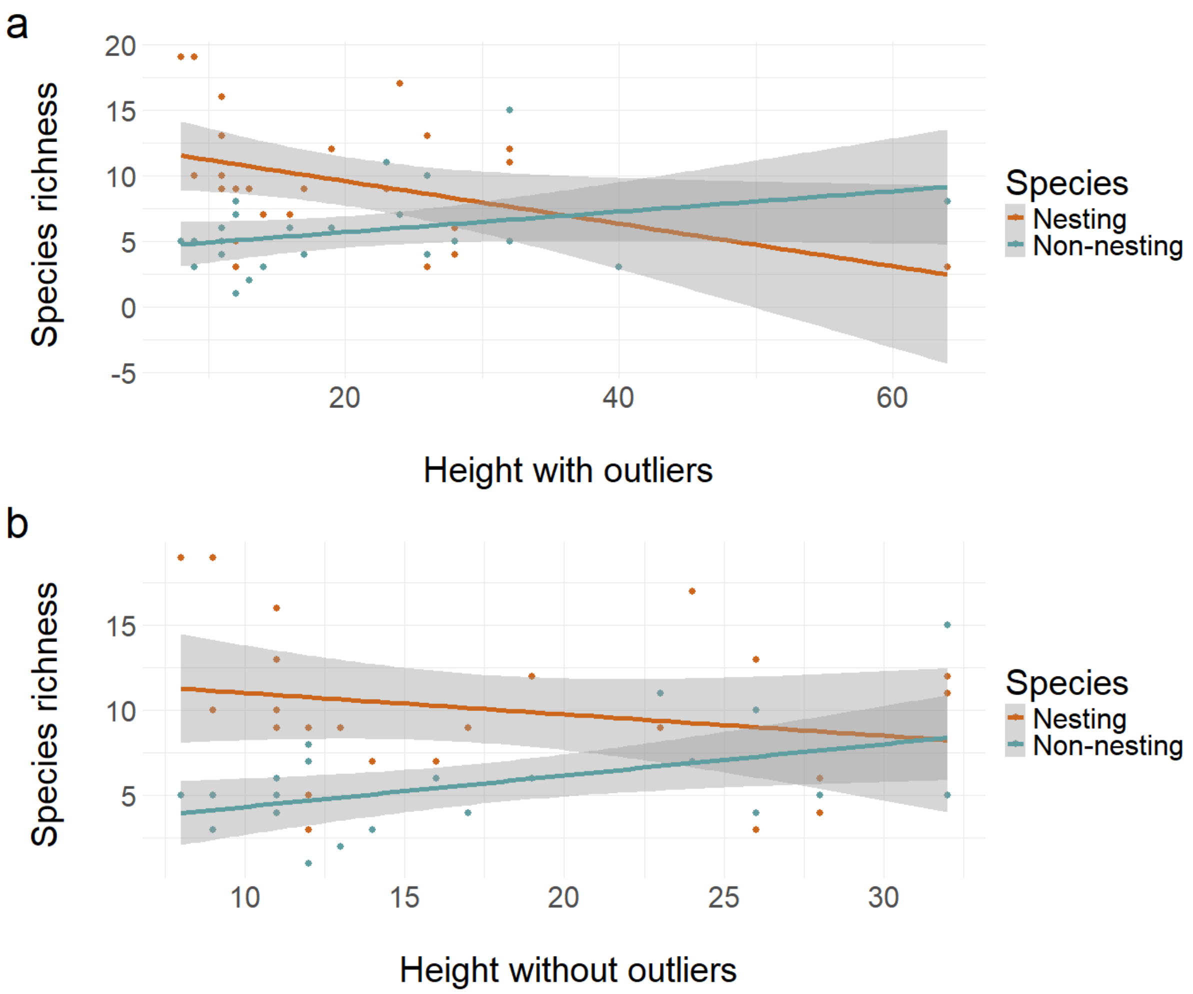

Fig. B.4. Height (x-axis) against species richness of non-nesting pollinators in blue and nesting pollinators in red (y-axis).(a) shows the trend with confidence intervals with the two outlier roofs that were over 35 meters. (b) shows the trend with confidence intervals without the outliers.

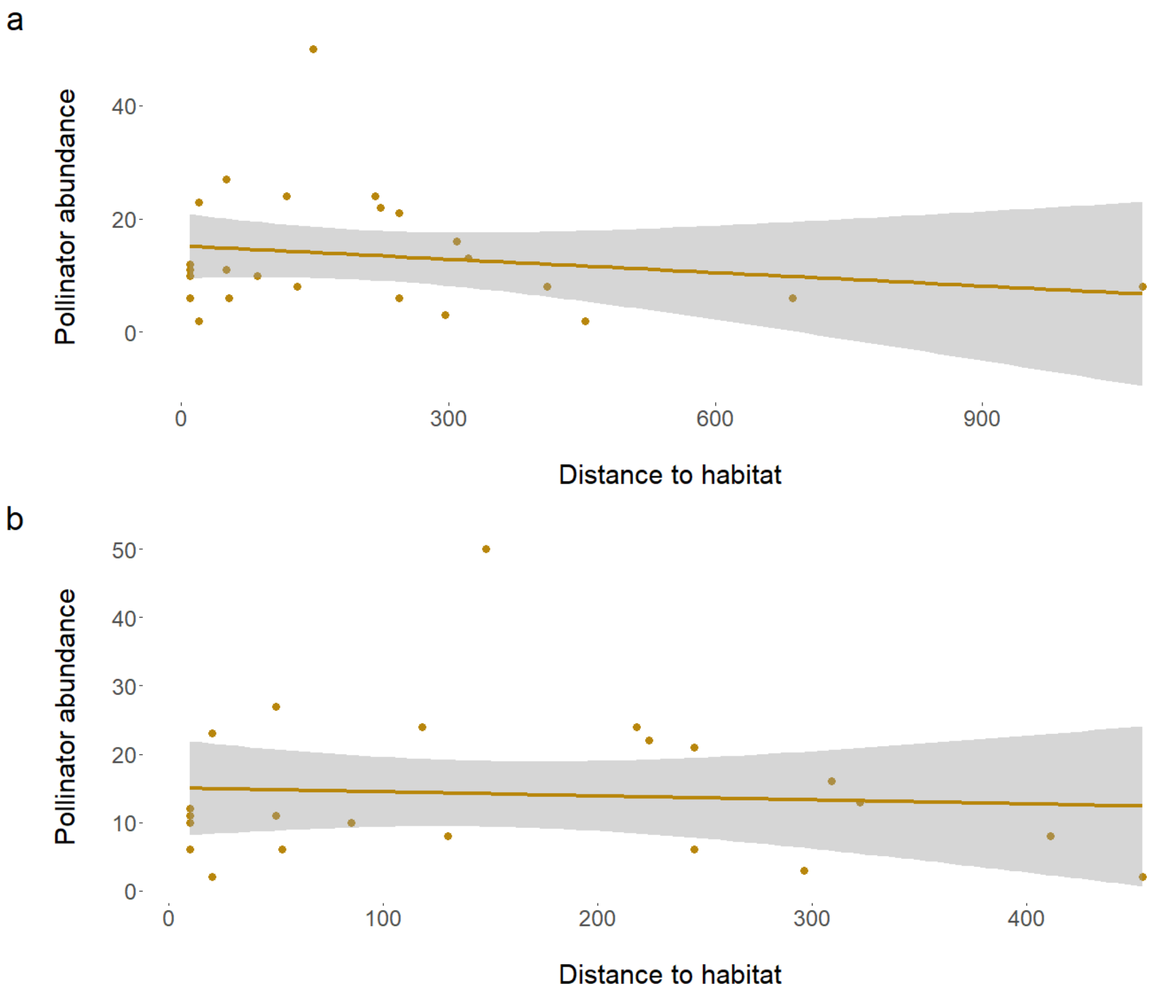

Fig. B.5. Distance to suitable habitat (x-axis) against abundance of non-nesting pollinators(y-axis).(a) shows the trend with confidence intervals with the two outlier roofs that were over 600 meters away. (b) shows the trend with confidence intervals without the outliers.
